# PAN-INDIA 1000 SARS-CoV-2 RNA Genome Sequencing Reveals Important Insights into the Outbreak

**DOI:** 10.1101/2020.08.03.233718

**Authors:** Arindam Maitra, Sunil Raghav, Ashwin Dalal, Farhan Ali, Vanessa Molin Paynter, Dhiraj Paul, Nidhan K Biswas, Arup Ghosh, Kunal Jani, Sreedhar Chinnaswamy, Sanghamitra Pati, Arvind Sahu, Debashis Mitra, Manoj Kumar Bhat, Satyajit Mayor, Apurva Sarin, The PAN-INDIA 1000 SARS-CoV-2 RNA Genome Sequencing Consortium, Yogesh S. Sauche, Aswin Sai Narain Seshasayee, Dasaradhi Palakodeti, Murali D. Bashyam, Ajay Parida, Saumitra Das

**Author notes:** Joint First Authors (Arindam Maitra, Sunil Raghav, Ashwin Dalal, Farhan Ali, Vanessa Molin Paynter, Dhiraj Paul). Corresponding Authors: Saumitra Das, Ajay Parida, Murali D. Bashyam, Dasaradhi Palakodeti, Aswin Sai Narain Seshasayee, Yogesh Sauche, Arindam Maitra. **PAN-INDIA 1000 SARS-CoV-2 RNA Genome Sequencing Consortium Team** **National Institute of Biomedical Genomics, Kalyani:** Arindam Maitra, Nidhan K. Biswas, Sreedhar Chinnaswamy, Shekhar Ghosh, Sumanta Sarkar, Subrata Patra, Rajiv Kumar Mondal, Trinath Ghosh, Arnab Ghosh, Shouvik Chakraborty, Saumitra Das **Institute of Life Sciences, Bhuabneswar:** Sunil Raghav, Arup Ghosh, Atimukta Jha, Swati Madhulika, Manashi Priyadarshini, Viplov K. Biswas, P. Sushree Shyamli, Bharati Singh, Neha Singh, Deepika Singh, Ankita Datey, Jyotirmayee Turuk, Jyotsnamayee Sabat, Debdutta Bhattacharya, Jaya Singh Kshatri, Rupesh Dash, Shantibhushan Senapati, Tushar K. Beuria, Rajeeb Swain, Soma Chattopadhyay, Gulam Hussain Syed, Punit Prasad and Ajay Parida **Centre for DNA Fingerprinting and Diagnostics, Hyderabad:** Ashwin Dalal, Murali Bashyam, Pratyusha Bala, Vinay Donipadi, Divya Vashisht, Debashis Mitra **Institute For Stem Cell Science and Regenerative Medicine-National Centre for Biological Sciences, Bengaluru:** Dasaradhi Palakodeti, Aswin Sai Narain Seshasayee, Uma Ramakrishnan, Shah-e-Jahan Gulzar, Varadharajan Sundaramurthy, Srikar Krishna, Vanessa Molin Paynter, Awadhesh Pandit, Farhan Ali, Mohak Sharda, Satyajit Mayor, Apurva Sarin **National Centre for Cell Science, Pune:** Dhiraj Paul, Kunal Jani, Janesh Kumar, Radha Chauhan, Vasudevan Seshadri, Girdhari Lal, Arvind Sahu, Yogesh S Shouche, Manoj Kumar Bhat **ICMR-National Institute of Cholera and Enteric Diseases, Kolkata:** Shanta Dutta, Mamta Chawla Sarkar, Ananya Chatterjee, Hasina Banu, Agniva Majumdar **Institute of Post Graduate Medical Education and Research, Kolkata:** Monimoy Banerjee, Raja Ray, Jayeeta Halder, Aritra Biswas **Translational Health Science and Technology Institute, Faridabad:** Guruprasad Medigeshi, Gagandeep Kang, Sharanabasava Patil, Anbalagan Ananthraj, Madhu Pareek, Imran Khan, ESIC Hospital and Medical College, Faridabad, Gurugram Civil Hospital, Gurugram and Palwal Civil Hospital, Palwal **Indian Institute of Science, Bengaluru:** Bharath K Sundararaj, Harsha Raheja, N. Srinivasan, Deepak K Saini, Amit Singh, K N Balaji, Umesh Varshney **Government Medical College, Aurangabad:** Jyoti Iravan, Dhaval Khatri, Maitrik Dave **All India Institute of Medical Sciences, Rishikesh:** Ravi Kant, Deepjyoti Kalita, Amit Mangla **Mahatma Gandhi Institute of Medical Sciences, Wardha:** Vijayshri Deotale, Rahul Narang, Deepashri Maraskolhe **Nizam’s Institute of Medical Sciences, Hyderabad:** K Manohar, Madhumohan Rao, Vijay Dharma Teja **Maulana Azad Medical College, Delhi:** Sonal Saxena, Vikas Manchanda, Oves Siddiqui **Regional Medical Research Center, Bhubaneswar:** Sanghamitra Pari, Jyotirmayee Turuk **Armed Forces Medical College, Pune:** Sourav Sen, Santosh Karade, KavitaBala Anand, Shelinder Pal Singh Shergill, Rajiv Mohan Gupta **Byramjee Jeejeebhoy Government Medical College, Pune:** Rajesh Karyakarte, Suvarna Joshi, Murlidhar Tambe.

## Abstract

The PAN-INDIA 1000 SARS-CoV-2 RNA Genome Sequencing Consortium has achieved its initial goal of completing the sequencing of 1000 SARS-CoV-2 genomes from nasopharyngeal and oropharyngeal swabs collected from individuals testing positive for COVID-19 by Real Time PCR. The samples were collected across 10 states covering different zones within India. Given the importance of this information for public health response initiatives investigating transmission of COVID-19, the sequence data is being released in GISAID database. This information will improve our understanding on how the virus is spreading, ultimately helping to interrupt the transmission chains, prevent new cases of infection, and provide impetus to research on intervention measures. This will also provide us with information on evolution of the virus, genetic predisposition (if any) and adaptation to human hosts.

One thousand and fifty two sequences were used for phylodynamic, temporal and geographic mutation patterns and haplotype network analyses. Initial results indicate that multiple lineages of SARS-CoV-2 are circulating in India, probably introduced by travel from Europe, USA and East Asia. A2a (20A/B/C) was found to be predominant, along with few parental haplotypes 19A/B. In particular, there is a predominance of the D614G mutation, which is found to be emerging in almost all regions of the country. Additionally, mutations in important regions of the viral genome with significant geographical clustering have also been observed. The temporal haplotype diversities landscape in each region appears to be similar pan India, with haplotype diversities peaking between March-May, while by June A2a (20A/B/C) emerged as the predominant one. Within haplotypes, different states appear to have different proportions. Temporal and geographic patterns in the sequences obtained reveal interesting clustering of mutations. Some mutations are present at particularly high frequencies in one state as compared to others. The negative estimate Tajimas D (D = −2.26817) is consistent with the rapid expansion of SARS-CoV-2 population in India. Detailed mutational analysis across India to understand the gradual emergence of mutants at different regions of the country and its possible implication will help in better disease management.

## Background

The ongoing pandemic of Severe Acute Respiratory Syndrome (SARS-CoV-2) has emerged as a global health problem reaching dimensions unparalleled in the history of humankind. Since its emergence in Wuhan, China in December 2019, it has spread rapidly to the rest of the world, with 17,354,751 people infected (as on August 1, 2020) and 674,291 deaths (1). Unlike SARS-CoV and MERS-CoV, the SARS-CoV-2 virus is characterised by an efficient person to person transmission, mostly via droplets, resulting in a basic reproductive number (R_0_) of 2.79 (1.5 and 6.68) (2) compared to R_0_ of 2·0–3·0, 0.9 and 1.5 for the SARS-CoV and the 1918 influenza pandemic, MERS-CoV, and the 2009 influenza pandemic respectively (3). Beginning from the first reported case in India in January 2020, the number of infections has risen to 1,695,988 including 36,511 deaths as on date (1, 4). Until date, 1,98,21,831 individuals in India have been tested for the infection (5).

Whole-genome sequencing of pathogens, especially viruses, is a powerful tool to generate rapid information on outbreaks, resulting in effective understanding of the introduction of the infection, dynamics of transmission, contact tracing networks and impact of informed outbreak control decisions (6–8). This will also provide us with information on evolution of the virus, genetic predisposition (if any) and adaptation to human hosts. In earlier outbreaks of the West African Ebola virus infection, rapid whole genome sequencing and analysis coupled with epidemiological information has been used effectively for public health decision making. In the ongoing pandemic, large scale whole genome sequencing initiatives in other countries are being used to establish the origins of the spreading infections as well as to determine how the virus has mutated during its transmission (9). Viral genome sequencing and epidemiological data have also been successfully used to estimate the impact of informed public health decisions on the spread of the virus and contact tracing of cluster cases (10). Hence, large scale generation of whole-genome sequence data on SARS-CoV-2 from clinical samples collected from multiple geographic locations and at different time points of the outbreak and rapid dissemination of the same in public databases like the Global Initiative on Sharing All Influenza Data (GISAID) (11), is expected to provide valuable information.

Since the initial sequence data from the first reported cases, there have been attempts by various medical and scientific research organisations to generate whole-genome sequence data from infected cases in India (12–15). Understanding the necessity of generation of large scale viral whole-genome sequences representing geographic locations all over the country and at different time points of the outbreak, a PAN-INDIA 1000 SARS-CoV-2 RNA Genome Sequencing Consortium of scientific research organisations and their collaborating clinical partners was formed under the aegis of Department of Biotechnology, Ministry of Science and Technology, Government of India (Figure 1). The Consortium members collected 1,062 clinical samples in the form of nasopharyngeal and oropharyngeal swabs from individuals testing positive for COVID-19 as per the guidelines of the Indian Council of Medical Research (ICMR), isolated and sequenced viral whole genomes. Collection and sequencing of additional samples is ongoing. The data is being jointly analysed along with available epidemiological information on these samples, and all efforts are being undertaken by the Consortium members to disseminate the sequence and related information in GISAID in real-time. It is expected that this large scale initiative will facilitate informed public health decisions to control the outbreak and provide major thrust in the development of intervention measures.

## Results

### Phylodynamic Analysis

Phylodynamic analysis using 1052 SARS-CoV-2 RNA sequences of DBT-Pan-India-Consortium was performed through Nextstrain/ncov pipeline, and time-tree was constructed after default quality control (Figure 2). 962 sequences out of 1052 sequences passed the stringent QC criteria in the Nextstrain/ncov pipeline. Sequences that are distributed over different regions of India (Northern India = 104, Western India = 201, Eastern India = 385, Southern India = 272) were classified into 5 different haplotypes, namely 19A (8%), 19B (5%), 20A (38%), 20B (48%) and 20C (0.5%) as described in Figure 2. In previously reported studies, 20A, 20B and 20C were reported as A2a haplotype. Further, 770 additional Indian SARS-CoV-2 sequences from GISAID (11) outside of DBT-Pan-India-Consortium were added to our dataset, and a phylodynamic time-tree was generated (Figure 3) which showed a similar pattern in viral haplotype distribution.

**Figure 01.**
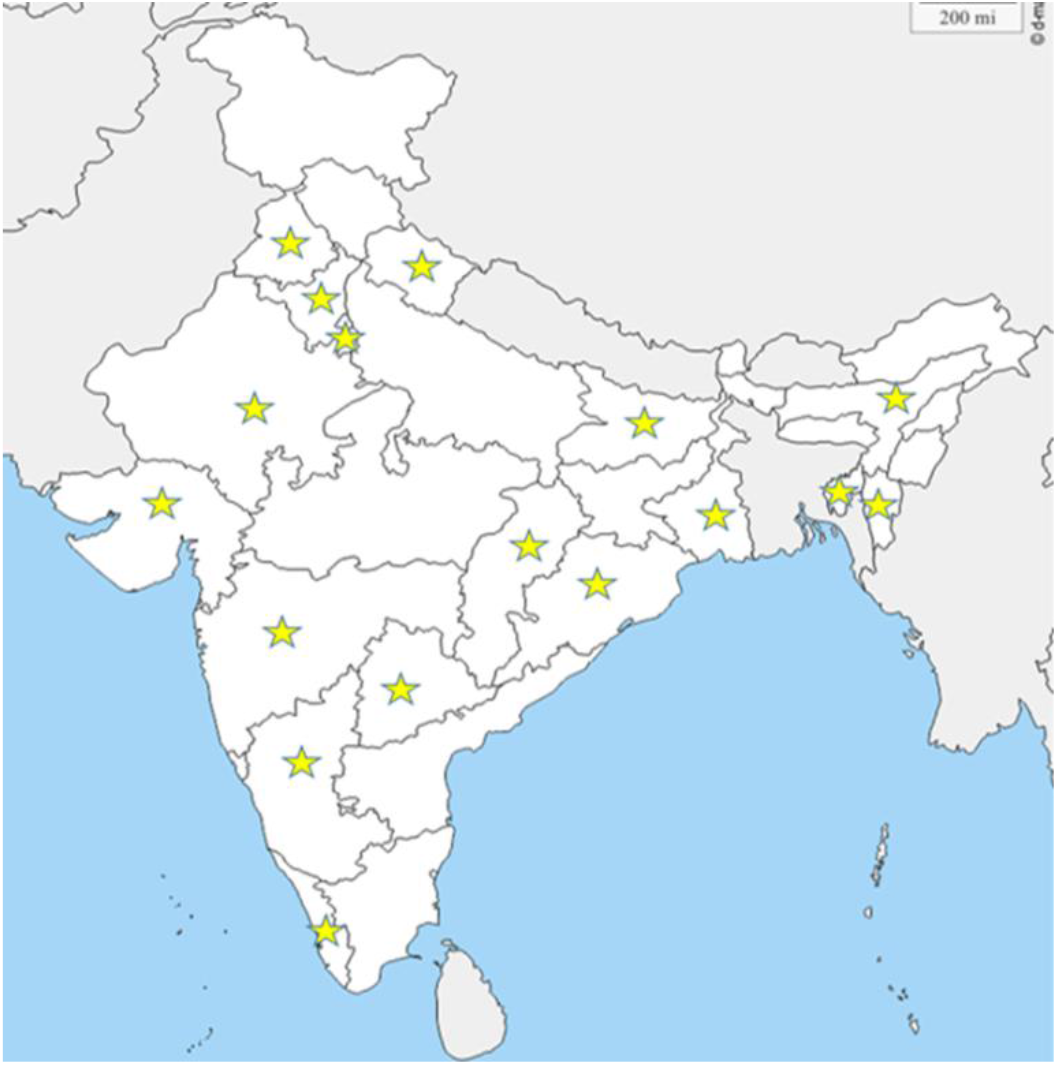
Regions of India covered by the PAN-INDIA 1000 SARS-CoV-2 RNA Genome Sequencing Consortium.

**Figure 02.**
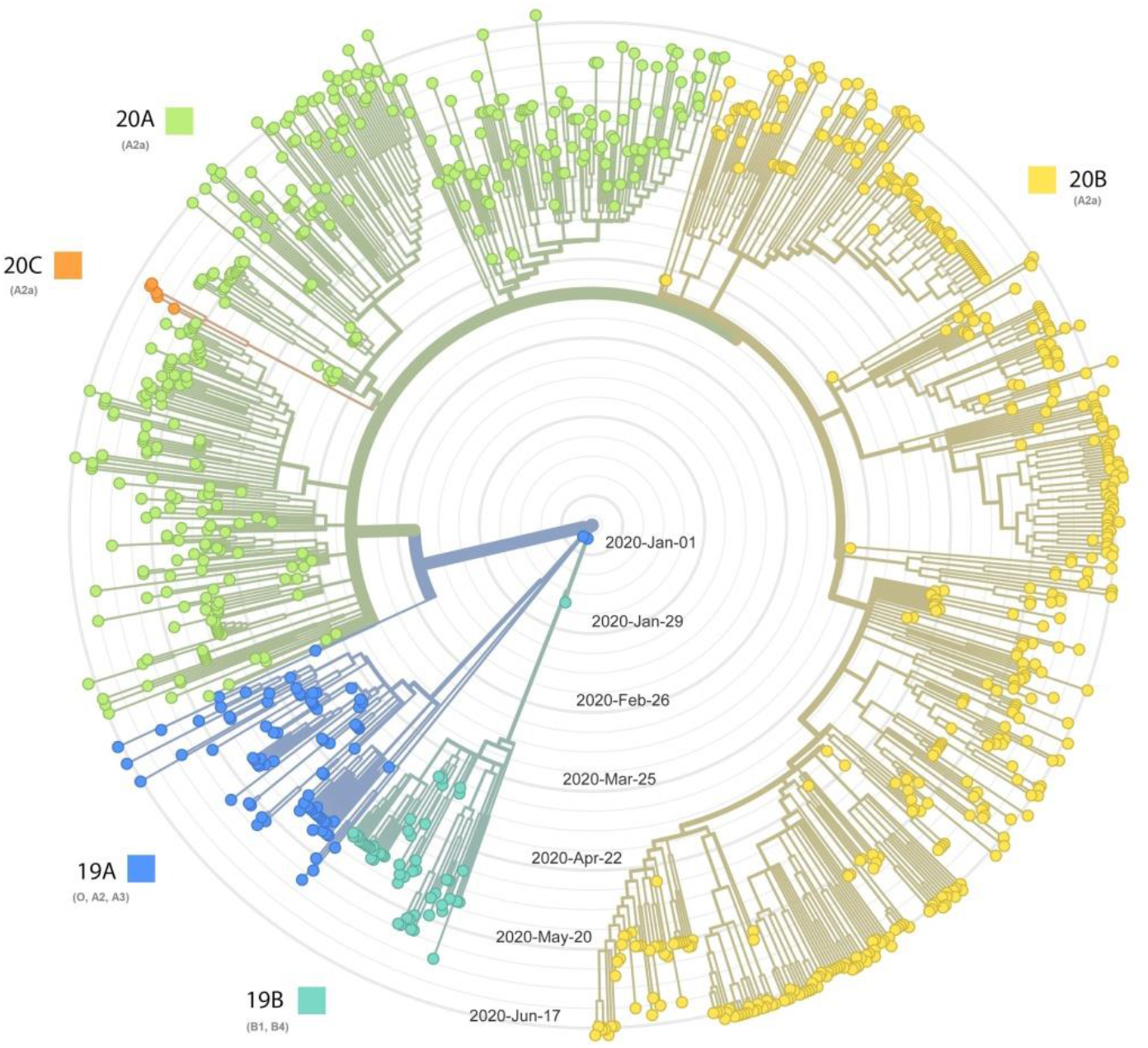
Radial phylogenetic timetree based on 962 SARS-CoV-2 RNA sequences. The concentric circles represent the sample collection date; the earlier date of sample collection is located close to the centre.

**Figure 03.**
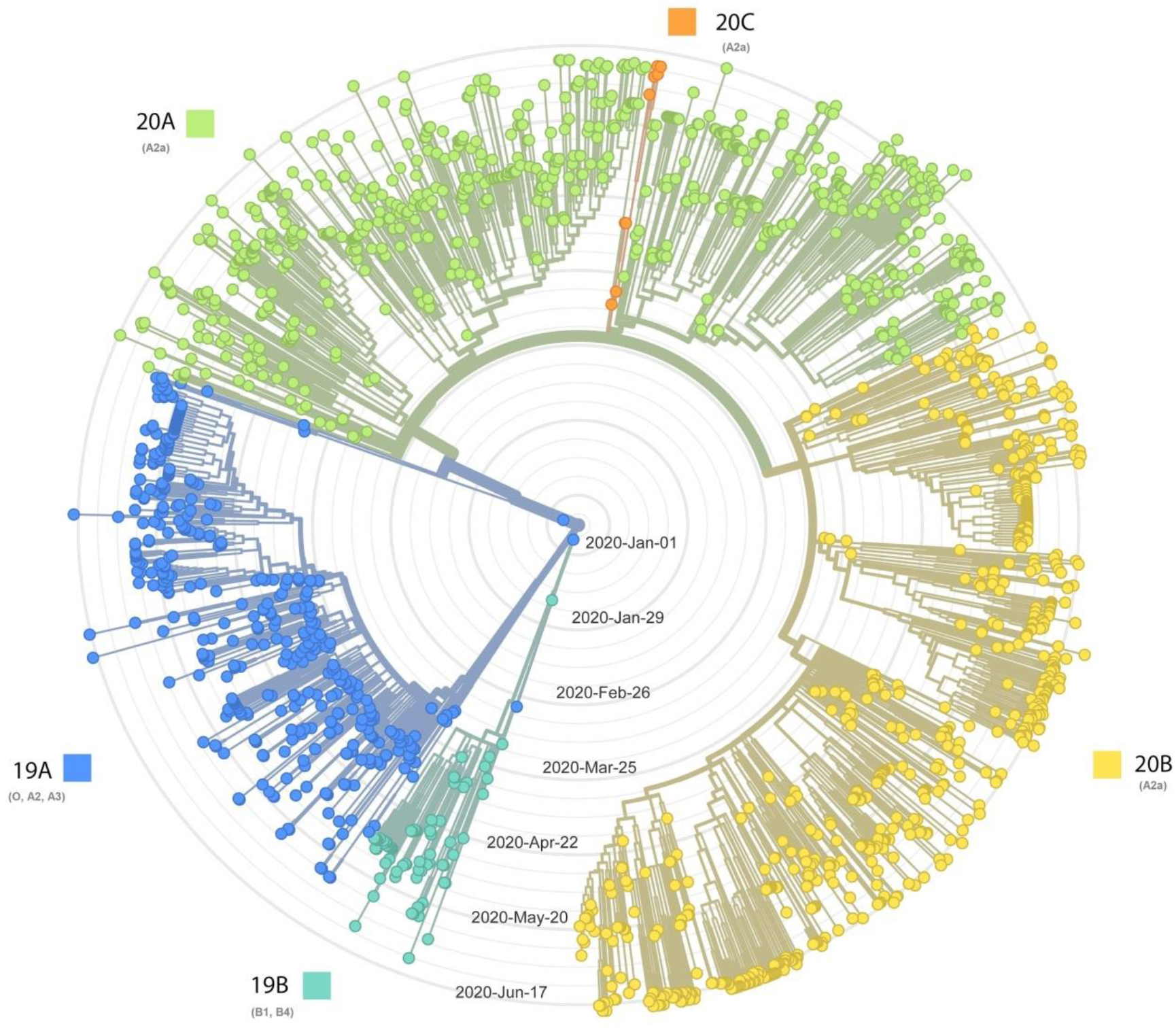
Radial phylogenetic timetree based on 1732 India-specific SARS-CoV-2 RNA sequences collected from this study and other studies.

SARS-CoV-2 RNA sequences generated by the Consortium were distributed in different states: 1) Northern India (Delhi, Uttar Pradesh, Haryana, Uttarakhand), 2) Eastern India (Odisha, West Bengal), 3) Western India (Maharashtra) and 4) Southern India (Telangana, Karnataka) as described in Figure 4.

**Figure 04.**
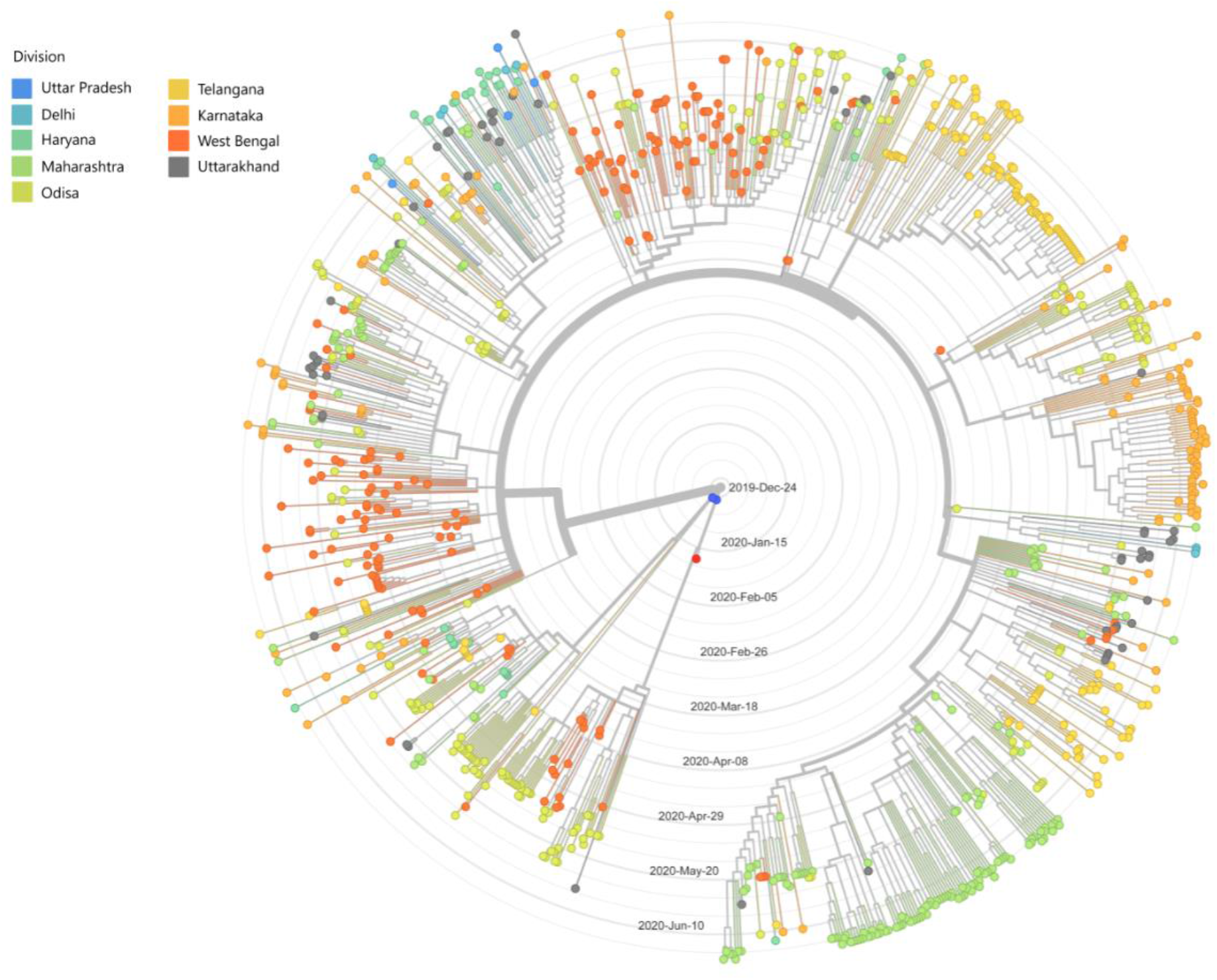
Distributions SARS-CoV-2 RNA sequences belonging to different states within India.

We have observed differential distributions of viral haplotypes in different parts of India as depicted by Figure 5 (state-wise) and Figure 6 (region wise). Viral clade 20A and 20B (previously known as A2a) dominated in all geographic regions of India. In Northern and Eastern India, the frequency of 20A (51.9% and 55.6%) was highest whereas 20B was found to be in higher frequencies in Western (70.6%) and Southern India (77.6%) compared to the rest. The frequency of ancestral 19A and 19B was higher in Northern (12.5%) and Eastern India (10.6%) as compared to Western (5%) and Southern India (5.1%). We have observed the highest frequency of 19B (13.2%) in Eastern India. We studied the differences in the haplotype distributions at pan India and regional levels over time. We find that although multiple haplotypes were introduced at the beginning of the outbreak (March-May 2020) in all regions, the predominant A2a haplotypes (20A, B and C) have overtaken others in June 2020.

**Figure 05.**
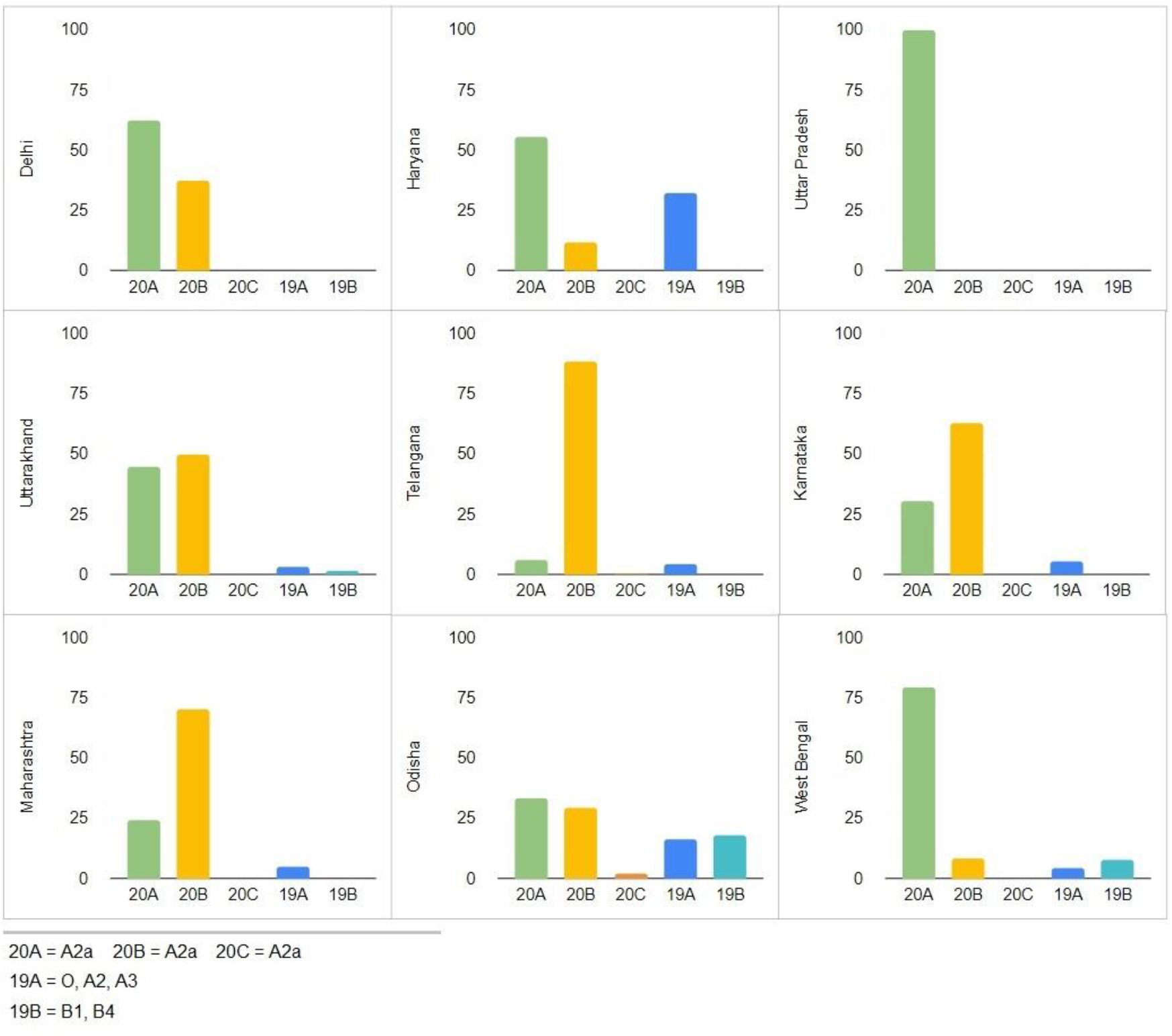
State-wise viral haplotype distributions.

**Figure 06.**
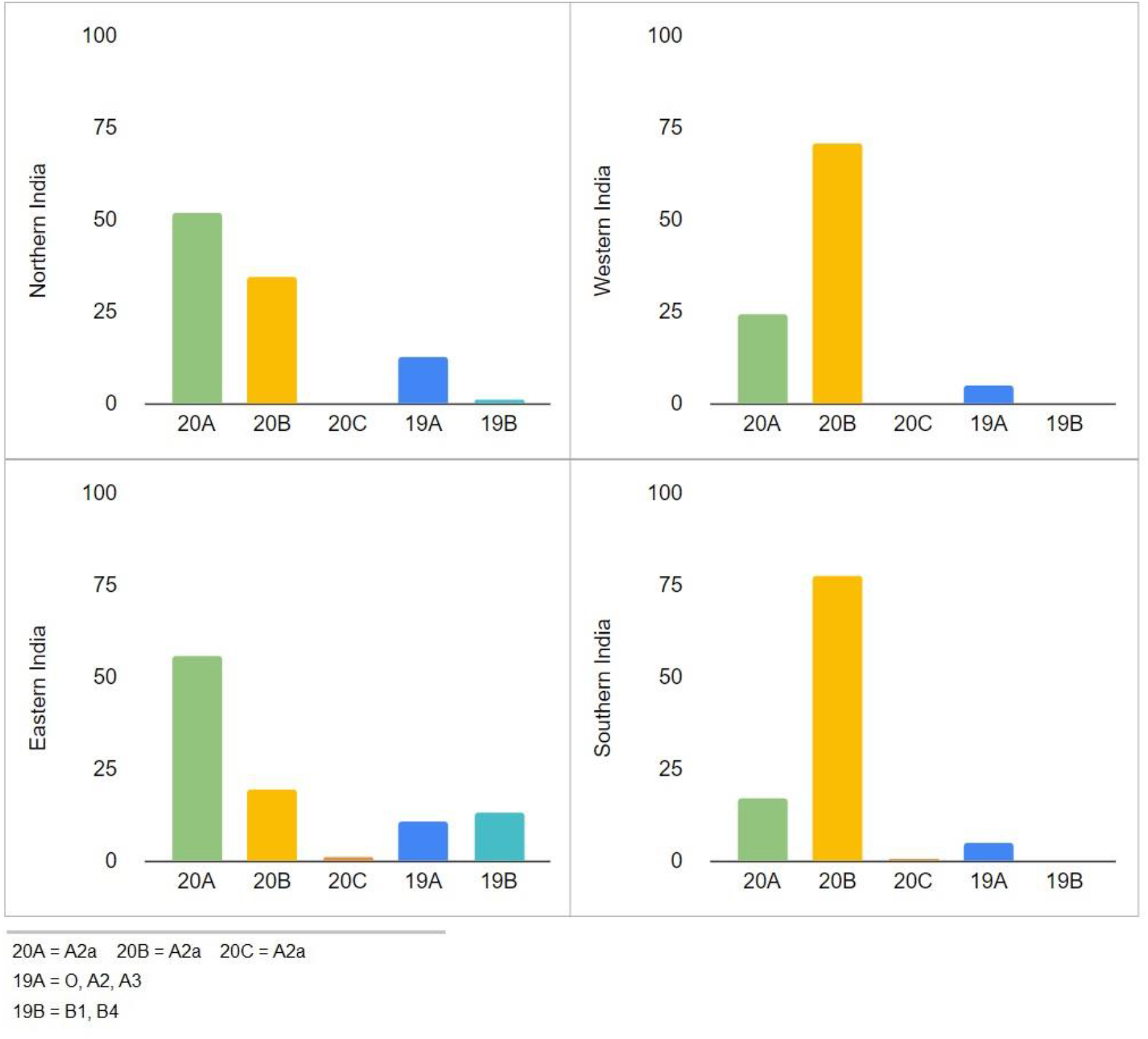
Region-wise viral haplotype distributions

Further, we have hypothesised that SARS-CoV-2 in India might be introduced from multiple foreign countries. To test our hypothesis, SARS-CoV-2 RNA sequences all over India were aligned to a subset of globally available sequences (~8000 random sequences representative of all months (Dec - July), all global region and all country locations) (Figure 7). Indian sequences that were classified into 19A and 19B haplotypes are majorly clustered with sequences from other Asian countries (Figure 7). In contrast, 20A, 20B and 20C sequences are clustered with European and American countries (Figure 7).

**Figure 07.**
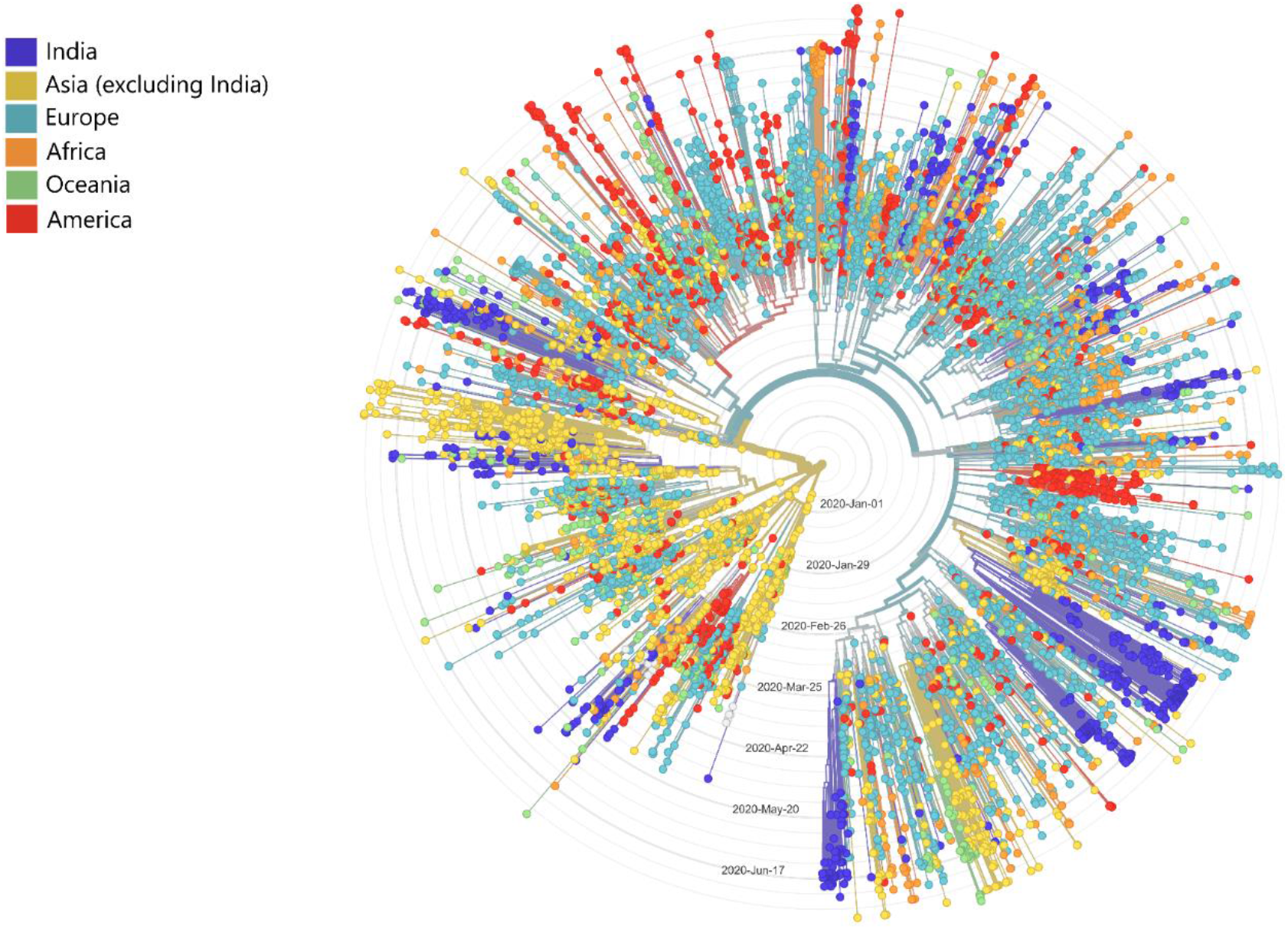
Phylodynamic timetree of Indian SARS-CoV-2 sequences with global subset consists of Asian, European, African, Oceanian and American countries.

In order to estimate possible SARS-CoV-2 introduction from foreign countries, we have utilised the country confidence information from phylodynamic timetree. Globally all sequences belonging to a particular clade were further divided into subgroups based on amino acid mutations (to estimate subgroup, the amino acid mutation is considered only if it is present in at least 10 samples for a particular clade). The country confidence information was extracted from the most recent common ancestors of each subgroup.

We have estimated SARS-CoV-2 introduction from multiple regions in India described in Figure 8 Further, we have visually validated the country confidence information from Nextstrain/auspice results for each geographic region of India (Figure 9, Figure 10, Figure 11 and Figure 12). Introduction of SARS-CoV-2 belonging to haplotype 19A and 19B mostly came from China and 20A, 20B and 20C from the United Kingdom, Italy and Saudi Arabia (Figure 8). We have noticed multiple introductions of the same haplotype in some geographic regions. Although we have found the same clade in different geographical regions of India, not all of them were introduced by the same foreign country. 20A haplotype of SARS-CoV-2 viruses came from Italy and Saudi Arabia in all Indian regions, but in the case of Eastern Indian, 20A came additionally from the United Kingdom and Switzerland. Similarly, 20B was introduced mostly from the United Kingdom in all regions, additionally in Western India from Brazil and in Southern India introduced from Italy and Greece. 19A was introduced from China in all regions. In contrast, 19B was introduced from Oman in North India and from China and Saudi Arabia in Eastern India.

**Figure 08.**
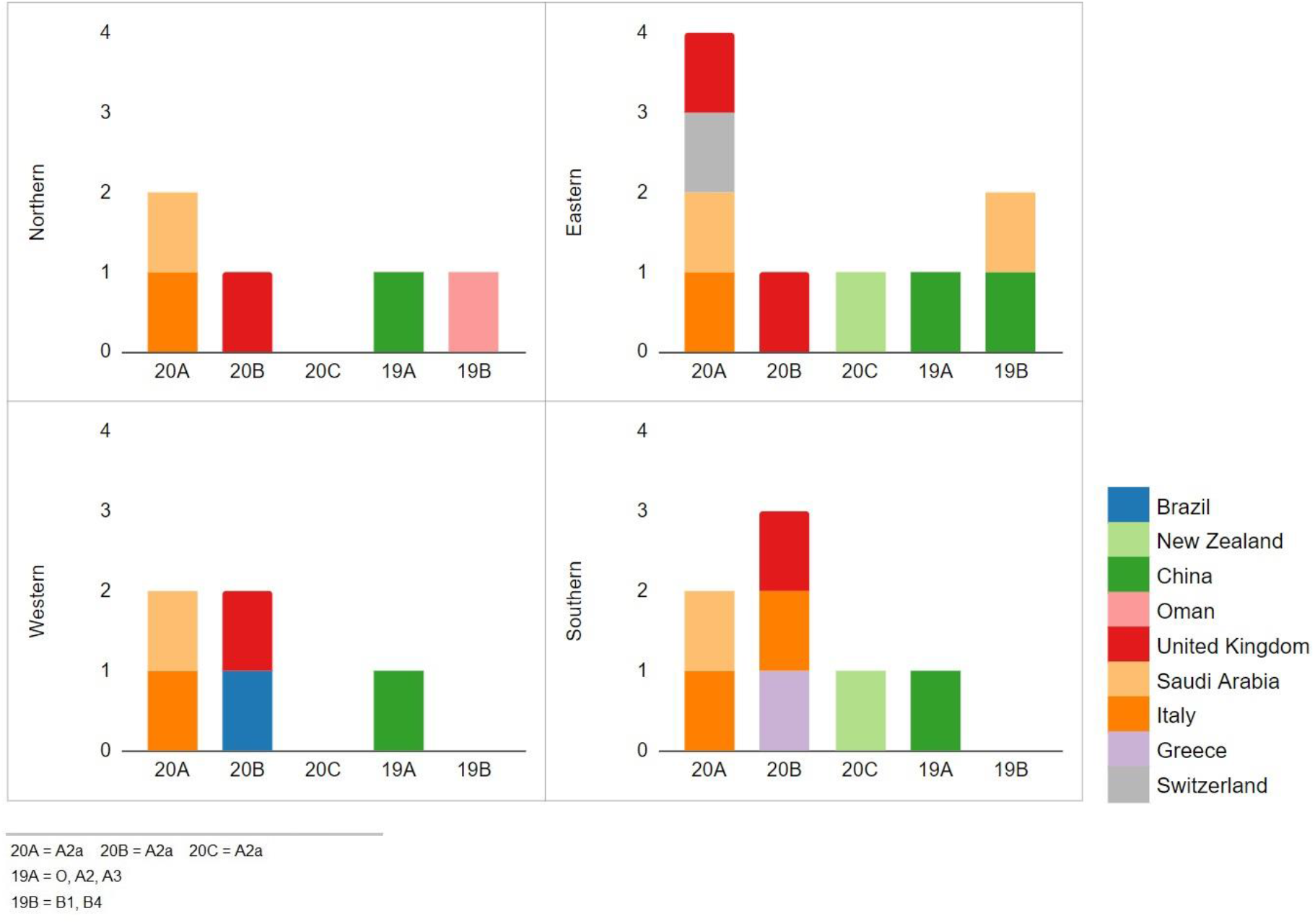
Introduction of SARS-CoV-2 haplotypes by foreign countries in different Indian regions. Foreign countries from which introductions were estimated are colour coded.

**Figure 09.**
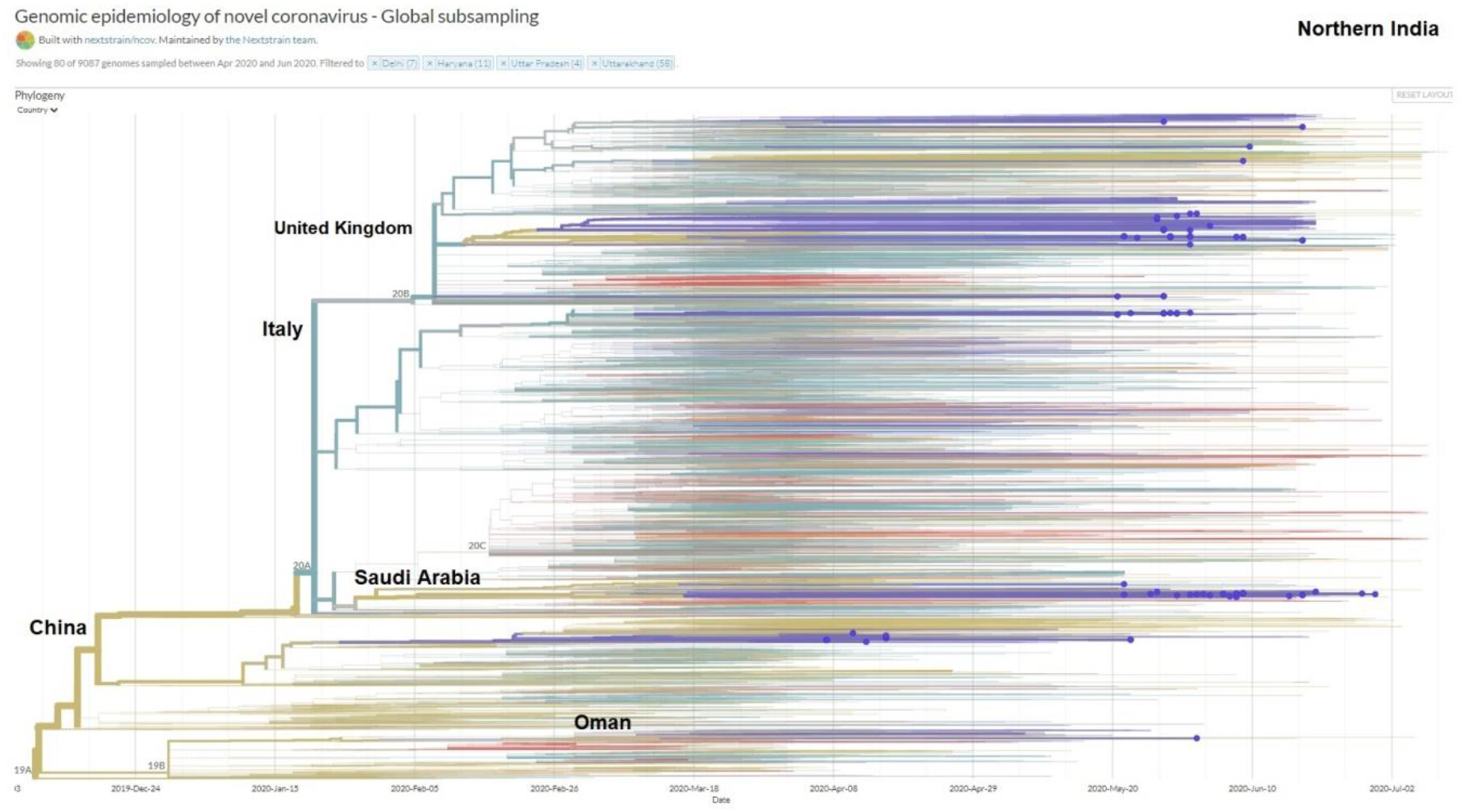
Northern Indian sequences in Timetree and estimated introduction by foreign countries.

**Figure 10.**
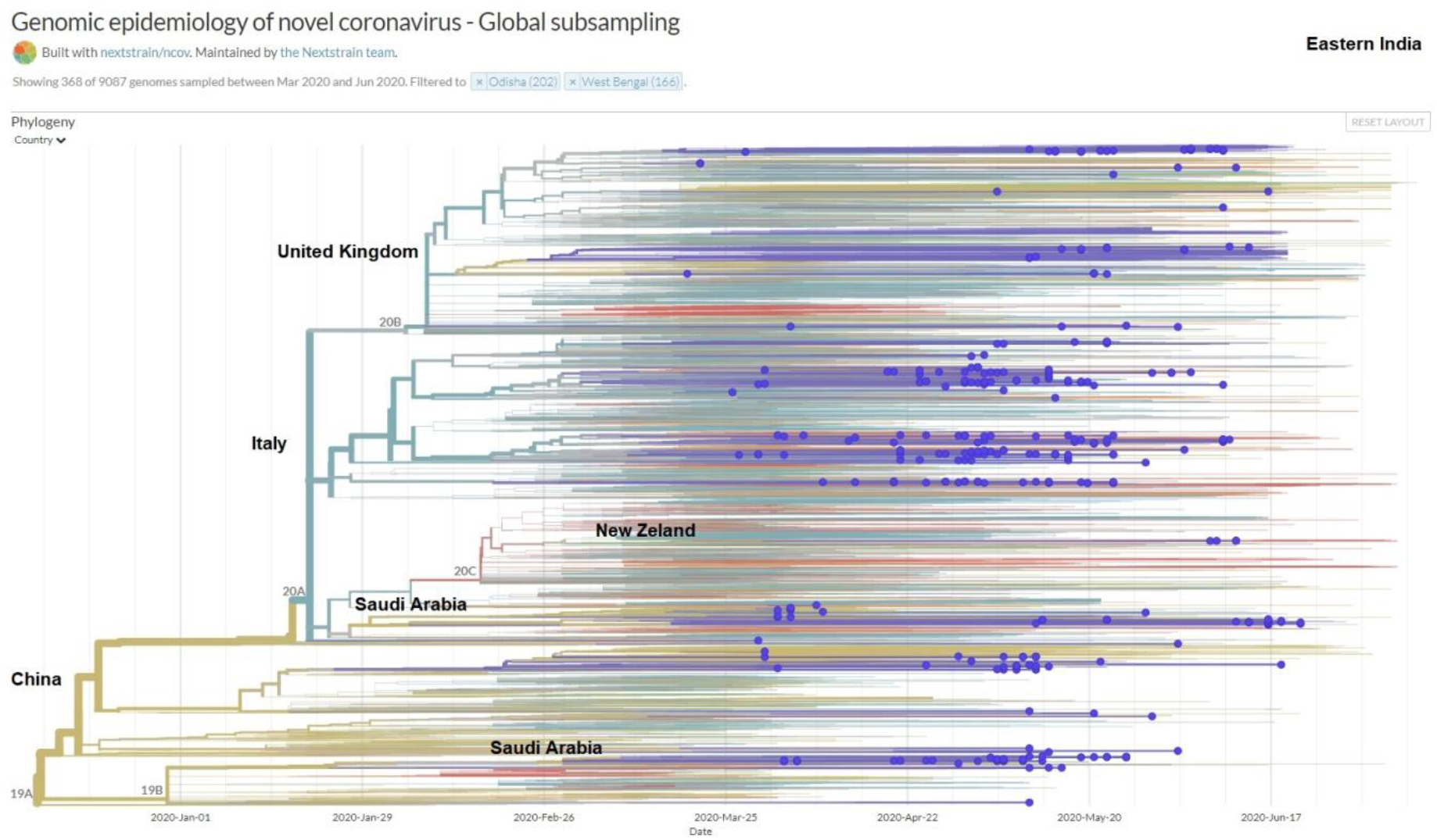
Eastern Indian sequences in Timetree and estimated introduction by foreign countries.

**Figure 11.**
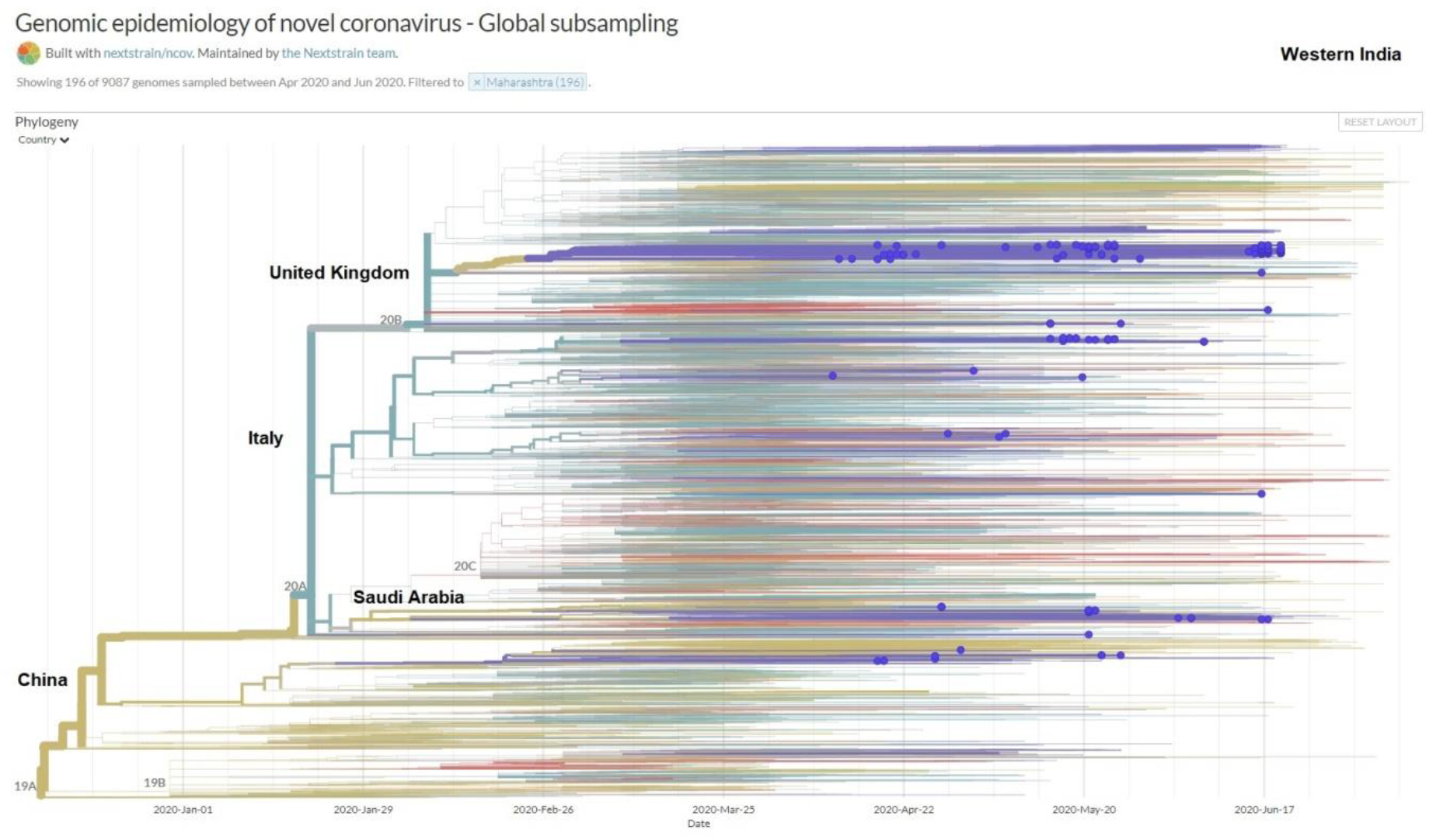
Western Indian sequences in Timetree and estimated introduction by foreign countries.

**Figure 12.**
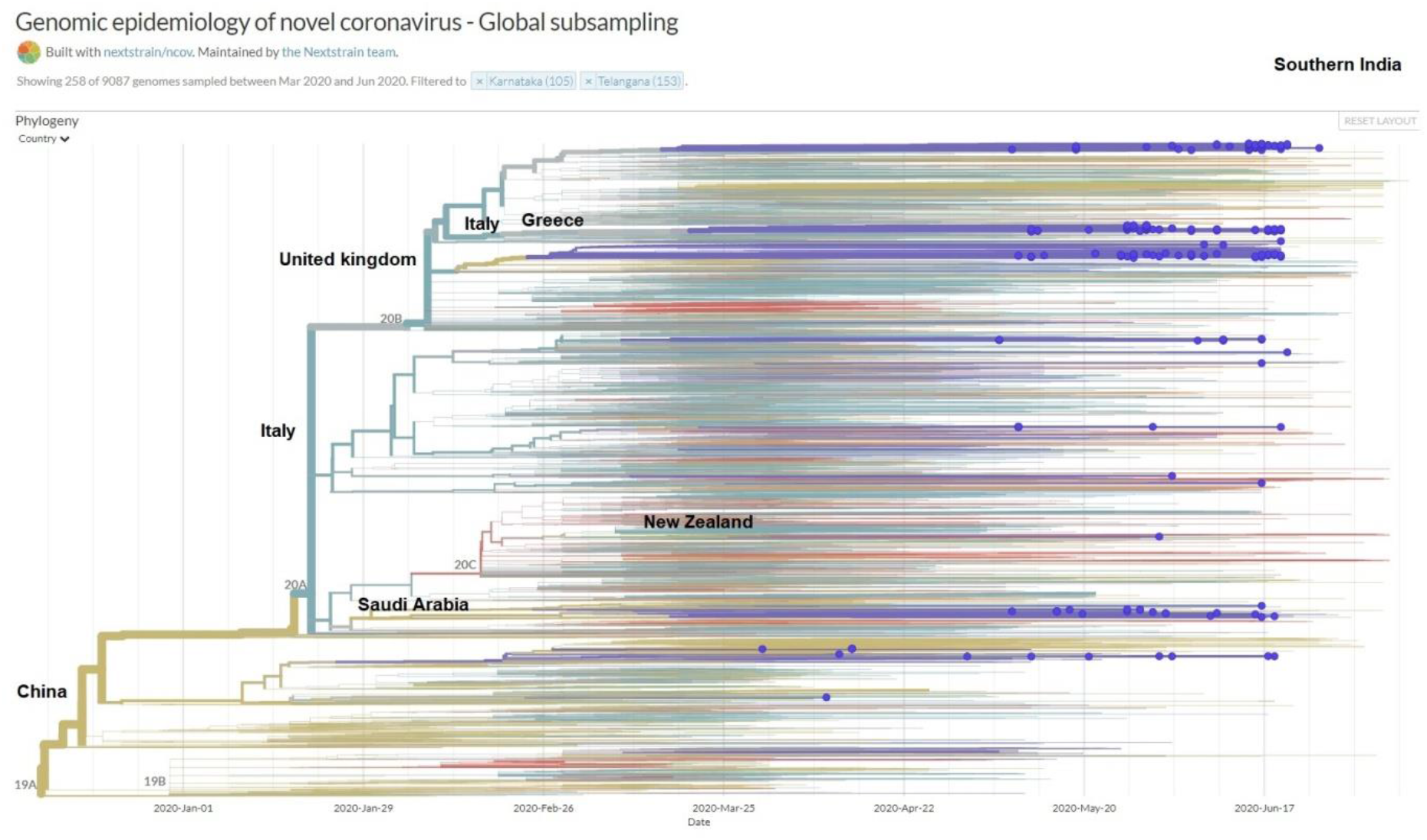
Southern Indian sequences in Timetree and estimated introduction by foreign countries.

### Mutation Analysis

To understand temporal and geographic patterns in the development of various genetic variants across India, we used a set of ~1,700 genomes combining those sequences by this consortium and others available in GISAID. We first defined sequence types based on the nucleotide sequence of 10 sites as defined by Guan et al. (16), and numbered according to the date of their first emergence globally, as described earlier for the CoVa pipeline (17). The most predominant sequence type among Indian samples was ST4 (Figure 13) which is characterized by two non-synonymous and 1 synonymous mutations (S D614G + RdRp P323L + nsp3 F106F). It represented 65 % of all samples. The second most predominant type was ST1 same as the reference type, and which accounted for 22 % of the dataset, followed by ST2 which is characterized by two mutations (ORF8 L84S + nsp4 S76S). The relative abundance of these types across states has changed over time (Figure 14). In most states, ST4 has increased at the expense of ST1, except in Delhi, where the distribution has shifted in the opposite direction.

**Figure 13.**
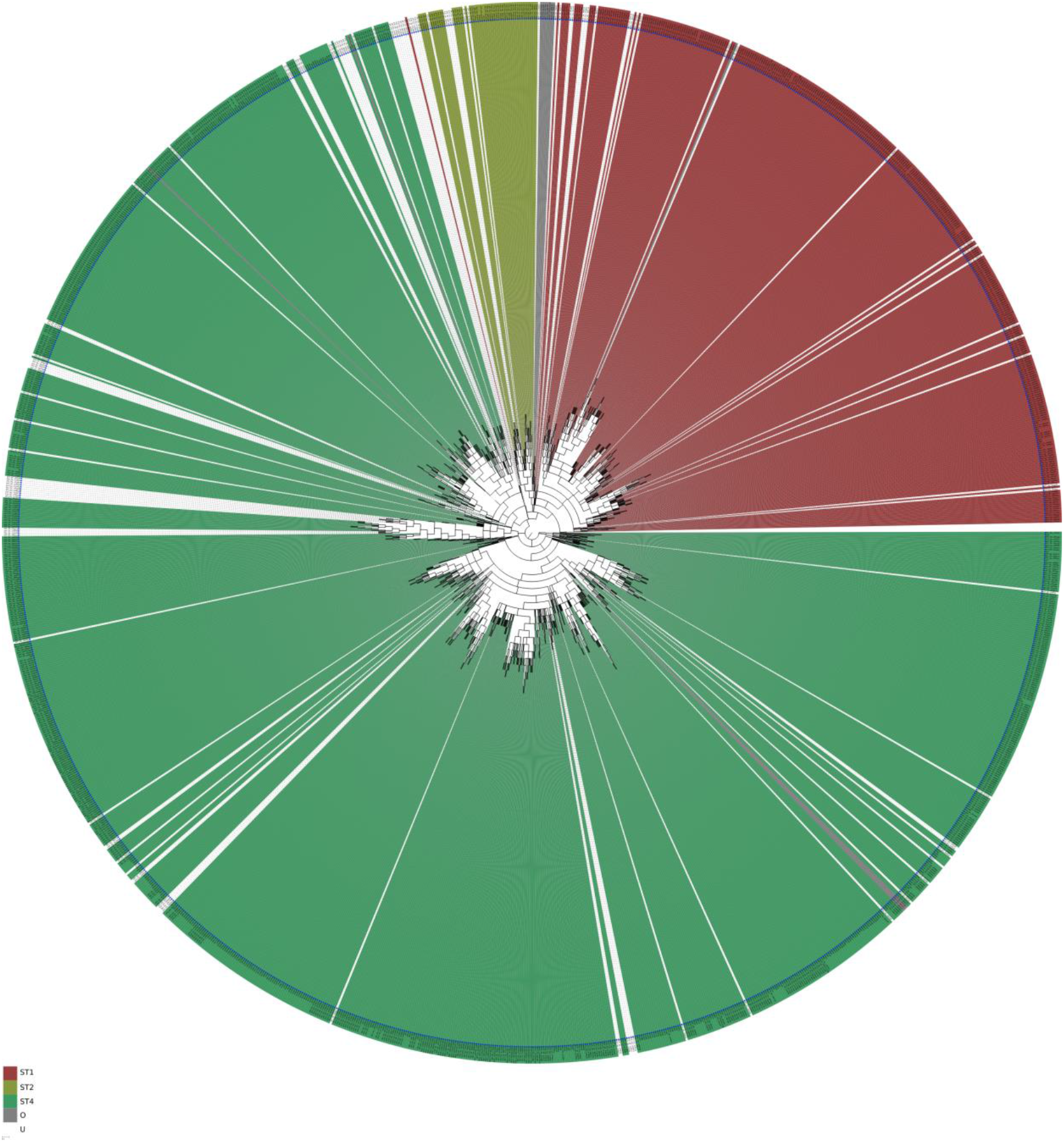
Phylogeny of 1630 SARS-CoV-2 genomes from India, depicting clustering and relative abundance of CoVa sequence types.

**Figure 14.**
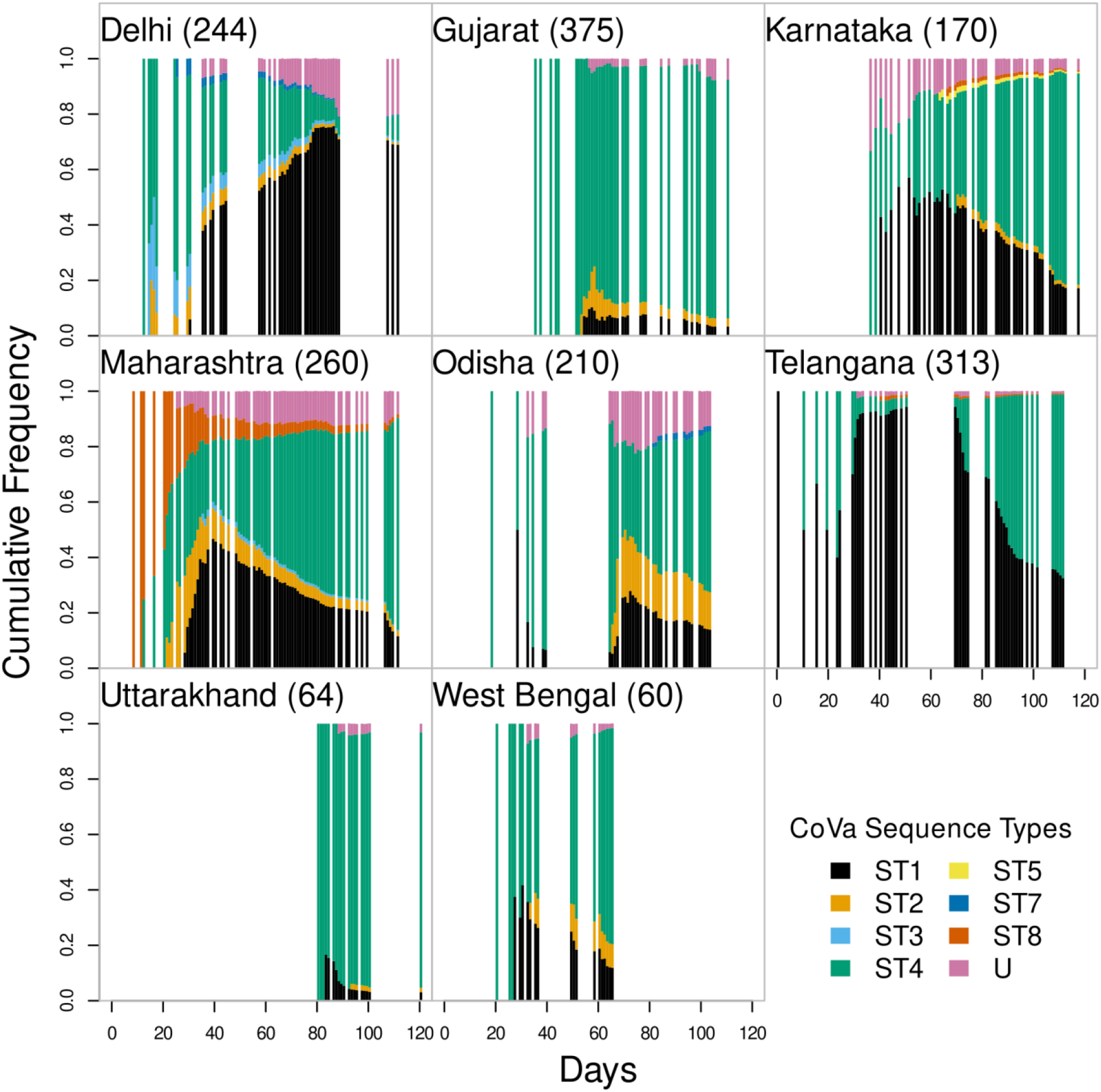
Changes in the relative abundance of sequence types across 8 Indian states.

The predominant type can also be observed by the frequency of its mutations. S D614G + RdRp P323L has attained frequencies above 75% in most states over time (Figure 15). In Delhi, however, the current majority is represented by 3 mutations (RdRp A97V + nsp3 T1198K + N P13L) which, considering the relative abundance of sequence types, have appeared not on the background of ST4 but ST1. Accordingly, this triad has decreased in frequency over time across other states. A sub-type seems to have attained high frequencies in multiple states; this one is characterized by two consecutive point mutations (N RG203KR) and appears to have originated on the background of type ST4.

**Figure 15.**
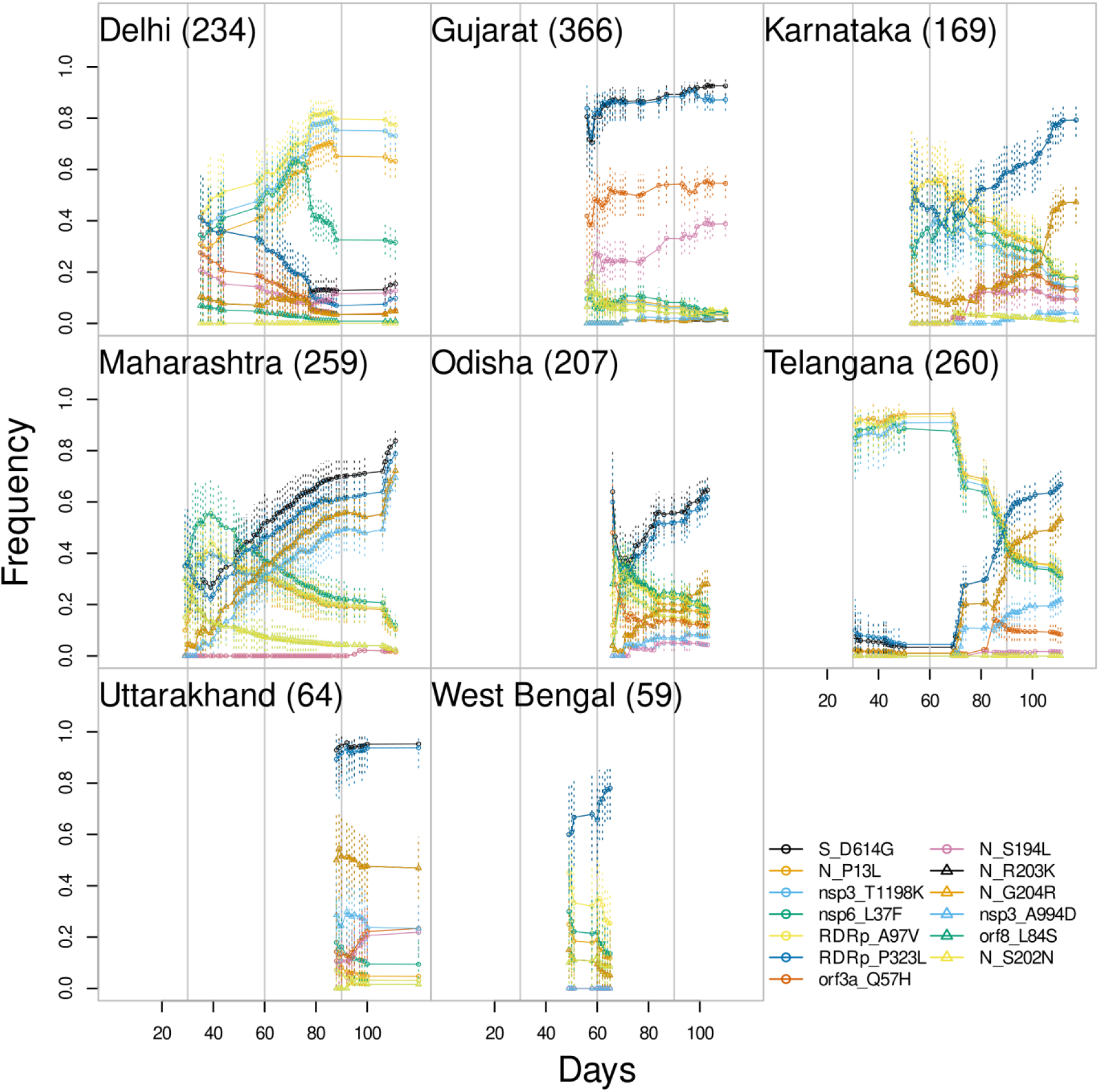
Frequency dynamics of major variants across 8 Indian states. Major variants were defined as those that were present in 1-3rd of the genomes in any state at least at one point of time.

Some mutations, other than the ones mentioned above, are present at particularly high frequencies in one state as compared to others. For example, ORF3a Q57H in Gujarat. One particular mutation that has gradually increased in frequency over time starting as a minority in a specific state is nsp3 A994D in Maharashtra, where it has reached a frequency above 50% while in other states, it is present at best at a frequency of 20%. The sharp peak in its curve towards the end corresponds to 86 sequences collected from Aurangabad in a span of 4 days. We note that such temporal and spatial sampling bias could affect other frequencies as well. However, the plot (Figure 15) is informative of temporal bias wherever the rise or decline of a mutation appears abrupt and is supported by a few data points, such bias is a likely explanation.

As seen above, frequency trajectories of many mutations appear correlated and indicate the presence of a few prevailing sub-types of the virus in the Indian population. Specifically, based on the average time-series correlation across states, two major clusters of mutations could be identified (Figure 16A). The two characteristic mutations of ST4 are strongly anti-correlated with a group of 4 mutations (nsp3 T1198K, RdRp A97V, N P13L, nsp6 L37F). On the other hand, they are positively correlated to a sub-cluster of 3 mutations (nsp3 A994D, N RG203KR). Some mutations might appear linked because of the artifacts of time-series correlation. If two mutations are not strongly monotonic in either direction, then they would appear correlated. Therefore, to directly assess the degree of co-occurrence of mutations, we performed clustering on the patterns of presence/absence of pairs of mutations across 1630 genomes. The results were largely similar (Figure 16B). However, the two pair of mutations, (ORF3a Q57H + NS194L) and (ORF8 L84S + N S202N) were separated into their own clusters.

**Figure 16.**
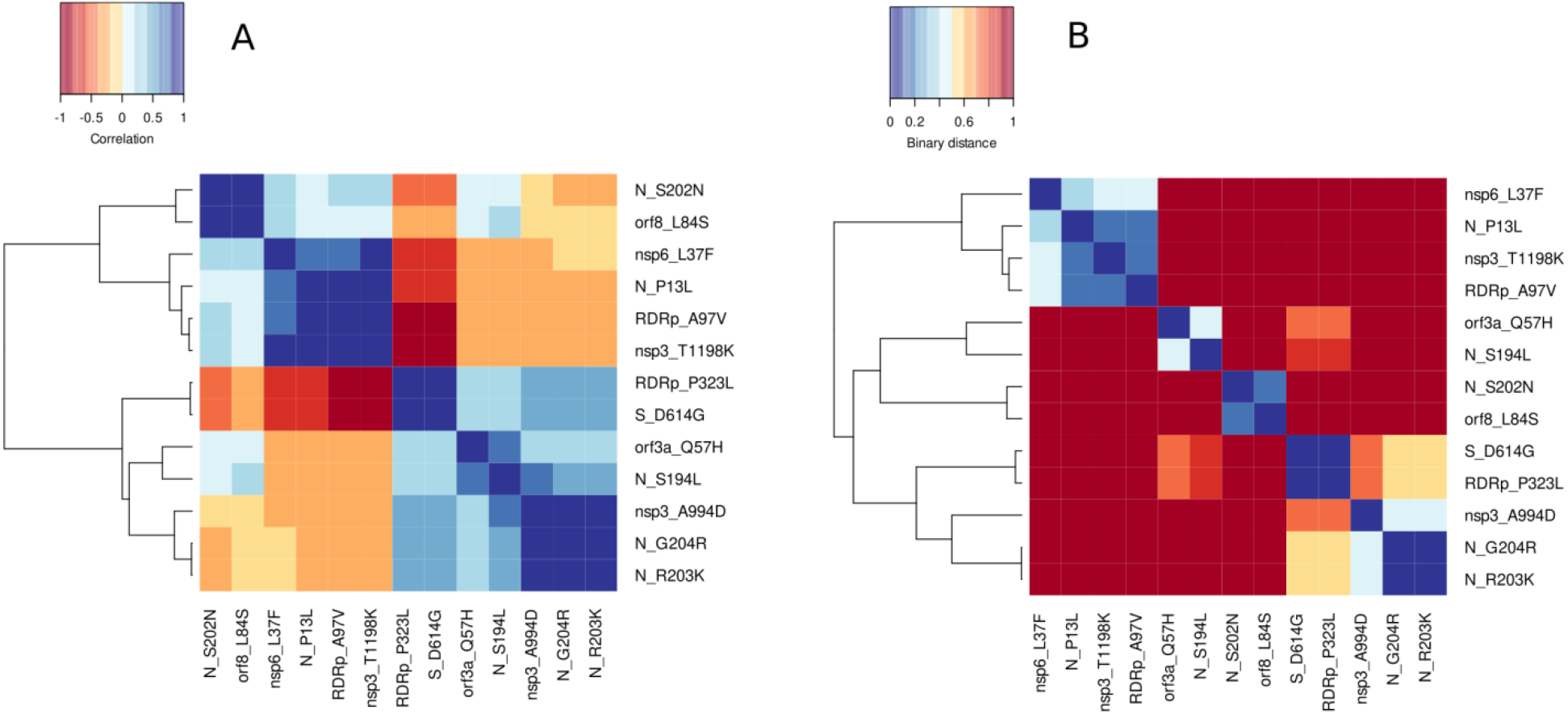
Clustering of mutations, **A**) Based on the frequency time-series correlation. **B**) Based on the pattern of co-occurrence across genomes.

### Haplotype network analysis

From a total of 1034 SARS-CoV-2 genome sequences we selected 717 high quality sequences representing the pathogen diversity in nine Indian states i.e. Delhi (n=8), Haryana (n=31),Karnataka (n=118), Maharashtra (n=115), Odisha(n=173), Telangana(n=24), Uttar Pradesh(n = 4), Uttarakhand (n=63) and West Bengal (n=181) were used for haplotype network re construction (Figure 17). From the haplotype network, we observed Maharashtra, Karnataka created three distinct haplotype nodes and sequences from Odisha, West Bengal and Uttarakhand sparse in different haplotype nodes. We also observed a haplotype node with the majority of the genomes from West Bengal, Odisha and a small percentage of the samples belonging to Uttarakhand. Geographically Odisha and West Bengal share borders, and the shared SARS-CoV-2 haplotypes might be because of the high interstate travelling. On the bottom right-side of the network we see a group of samples from Maharashtra, Delhi, Haryana and Uttarakhand grouped together with 2-4 single nucleotide variants (SNVs), suggesting the infection might have spread in a short duration of time. On the left of the network, there is a portion of the samples from Haryana and Karnataka sharing the same parent haplotype, representing possible transmission by migration. The negative estimate Tajima’s D (D = −2.26817, AMOVA p(D<=−2.26817) = 0.00281) (18) is consistent with the rapid expansion of SARS-CoV-2 population in India.

**Figure 17:**
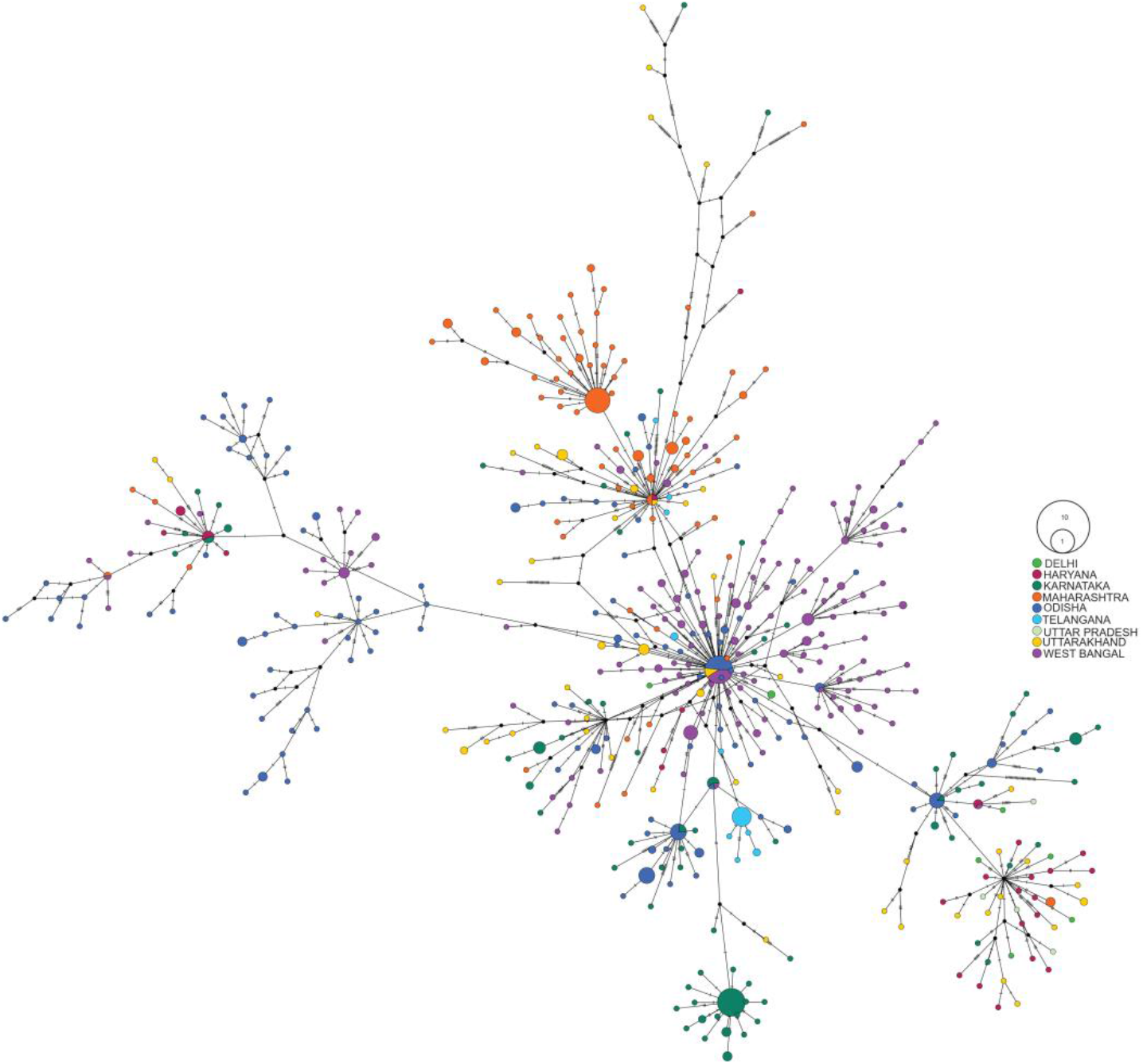
Haplotype network of 717 SARS-CoV-2 genomes (Delhi (n=8), Haryana (n=31), Karnataka (n=118), Maharashtra (n=115), Odisha (n=173), Telangana (n=24), Uttar Pradesh (n = 4), Uttarakhand (n=63), West Bengal (n=181)) from DBT’s PAN-India 1000 SARS-CoV2 RNA genome sequencing consortium. The circle size is proportional to the number of samples belonging to the haplogroup.

## Discussion

Our analysis shows the presence of multiple lineages of SARS-CoV-2 in India in different geographic regions. 20A and 20B together (belonging to the former haplotype of A2a) are the predominant haplotypes at pan India level and in each region. Interestingly, the haplotypes were differentially distributed in different regions. While the 20A were most abundant in Northern and Eastern India, 20B was found to be the most abundant haplotype in Western and Southern India. The ancestral haplotypes of 19A and 19B were mostly found in Northern and Eastern India, with 19B being the most abundant in the latter region. Our data shows interesting temporal changes in haplotype diversity of SARS-CoV-2 in India. While multiple haplotypes were introduced during the early part of the outbreak in March-May 2020, the 20A, B and C haplotypes (A2a) became the predominant haplotypes in all regions by June 2020. This is consistent with the analysis of global data (12) and the reported enhanced efficiency of transmission ability of the A2a haplotype and its association with high viral load (13, 14).

Analysis of probable country of origin of these SARS-CoV-2 sequences in India revealed that they had been probably introduced by travel from multiple countries across the globe. 20A, B and C haplotypes were introduced from multiple countries in Europe and also American continents. Interestingly, 20A alone is predicted to have been introduced by travel from Italy, Saudi Arabia, United Kingdom and Switzerland. Similarly, 20B was introduced from the United Kingdom, Brazil, Italy and Greece. In contrast, 19A was introduced from China alone while 19B was introduced by travel from China, Oman and Saudi Arabia.

The most predominant SARS-CoV-2 lineage across India is the one with D614G variant of the Spike protein. Across most states, it’s relative abundance has increased over time to 70% and above, except in Delhi, where an earlier lineage, ST1 (known as type L in GISAID) is the predominant type present at a similar proportion. We found three characteristic mutations for this lineage of type ST1 (nsp3 T1198K, RdRp A97V, N P13L). The frequency time-series of these two lineages across states was strongly anticorrelated. Based on the co-occurrence patterns and frequency dynamics, we identified a sublineage of D614G, characterized by two adjacent mutations in the nucleocapsid protein (N RG203KR), in many states. In Maharashtra, it was the predominant lineage and was strongly correlated to another mutation nsp3 A994D. This particular mutation had attained frequencies above 50% in Maharashtra while its end point frequency elsewhere was no greater than 20%. Further work would be required to understand the effect, if any, of these mutations on the infectivity or virulence of the virus.

An interesting set of mutations which has enriched during the course of the outbreak in India has been the emergence of the RG203KR in the N gene. Since its detection in few sequences obtained from individuals in April 2020 (15), this mutation has been found to gain abundance, especially in the West, South and North region but not in the East in our sequence data. Further analysis and studies are required to be undertaken to understand the implication of this observation, since this mutation is expected to alter potential miRNA binding sites and cause changes in the structure of the nucleocapsid.

Our analysis indicates that the haplotype diversities pan India and in each region continued to increase until May 2020, subsequent to which it reduced drastically with the emergence of the A2a haplotypes which has overtaken other lineages by June 2020. Further data is required to understand whether this observation might also reflect no new introductions of the virus in the country subsequent to May 2020 since India implemented a national lockdown between April to May. Such interpretations might enable improved understanding of such informed public health decisions. In recent times the number of COVID-19 occurrences in India has increased drastically. Although most of the states have their own strategic lockdown devised to control the outbreak, it will be more efficient if we can incorporate the geographical transmission pattern information in the planning of such strategies. In the current haplotype network, we have tried to explore the transmission of the infection among different states of India. It is necessary to incorporate more genomic datasets to draw a clearer picture.

## Methods

### Viral Genome Sequencing and analysis

RNA isolated from nasopharyngeal and oropharyngeal swabs was used to prepare genome sequencing library using the following kits as per manufacturer’s instructions; (i) TruSeq Stranded Total RNA Library Preparation Kit (Illumina Inc, USA) for shotgun metagenomics RNA sequencing, or (ii) QIAseq SARS-CoV-2 Primer Panel (Qiagen GmbH, Germany) for amplified of viral genome sequencing or (iii) Maxima H Minus Double-Stranded cDNA Synthesis Kit (ThermoFisher Scientific, USA), Nextera Flex Enrichment Kit with Respiratory Virus Oligo Panel (Illumina Inc, USA) for viral RNA capture and sequencing. All the sequencing libraries were checked using high sensitivity D1000 ScreenTape in 2200 TapeStation system (Agilent Technologies, USA) and quantified by Real-Time PCR using Library Quantitation Kit (Kapa Biosystems, USA). Next-Generation Sequencing were carried out using MiSeq Reagent Kit v3 or MiSeq Reagent Kit v2 Micro or Miseq reagent Kit v2 Nano in Miseq system (Illumina Inc, USA) or using NovaSeq 6000 SP Reagent Kit (Illumina Inc, USA). All of these libraries were prepared for 2×100 bp sequencing reads or 2×150 bp sequencing reads. For shotgun RNA sequencing data and captured viral RNA sequencing data, sequencing reads were mapped to reference viral genome sequence and consensus sequence for each sample was built using Dragen RNA pathogen detection software (version 9) in BaseSpace (Illumina Inc, USA). For amplified whole-genome sequencing, the viral sequences were assembled, and a consensus sequence for each sample was generated using CLC Genomics Workbench v20.0.3 (Qiagen GmbH, Germany). In both cases, the SARS-CoV-2 isolate Wuhan-Hu-1 (Accession NC_045512.2) was used as the reference genome.

### Phylodynamic Analysis

SARS-CoV-2 RNA sequences (N=1052) generated by DBT-all-India-Consortium representing major geogrphic regions in India were combined. To analyze the data, community standard Nextstrain/ncov pipeline was used. Nextstrain/ncov pipeline (19) includes quality control, alignment, phylogenetic inference, temporal dating of ancestral nodes and discrete trait geographic reconstruction which utilises MAFFT (20), augur and auspice tools.

Prior to analysis, preliminary quality control as sequence duplicate removal from multifasta and fixing of metadata information was performed. Sequences that did not met minimum length criteria, maximum ‘N’ content were removed. Over a period of time, Nextstrain curates extremely divergent samples in their GitHub repository, which were also removed by the pipeline from the dataset before alignment. 1010 sequences that passed QC criteria were finally aligned using MAFFT. Post alignment 46 sequences were removed, which includes clustered mutations and divergent samples. The timescale and branch lengths of phylogenetic tree were estimated using the IQ-TREE method (21) considering Wuhan/WH01/2019 as ancestral. Viral clades could be assigned to 962 sequences. Further, frequency-trajectories of mutations, genotypes and haplotypes (clades) were estimated by Augur. Finally, the phylogenetic timetree was constructed by auspice tool.

### Transmission profiles of SARS-CoV-2 sequences in India

In order to get an estimation of viral introduction from a foreign country to a geographical region in India, we prepared a global subset of 9201 sequences (Including all DBT-all-India-Consortium sequences). Around eight thousand foreign sequences were randomly subsampled from GISAID using Nextstrain subsample module, which represented all months (Dec - July), all global region and all country locations. A maximum of 75 sequences per month per country were taken during subsampling. Phylodynamic analysis with Indian SARS-CoV-2 sequences with global subset was performed as was described in the previous section.

Sequences assigned to a particular clade were further divided into subgroups based on amino acid mutations. Only those amino acid mutations that present in at least 10 samples in a particular clade were considered while performing subgrouping. The most recent common ancestors (MRCA) for each subgroup were determined using the ‘Phylo’ module of Biopython (22) from the phylogenetic tree. The country confidence information of MRCAs for each subgroup was extracted and curated from auspice timetree using python scripts. Further, we have visually validated the introduction analysis results from the auspice generated web-portal.

### Mutation Analysis

#### 1. Data description

1586 SARS-CoV-2 genome assemblies were available from India on GISAID as of July 18, 2020. Additionally, 631 assemblies, which were not yet available publicly, were directly provided by NIBMG and NCCS. A total of 1058 genomes were sequenced by the consortium till that date. Out of these 2217 genomes, 1928 had information on the collection date as well as on the state location. 8 Indian states (Karnataka, Maharashtra, Telangana, Odisha, West Bengal, Uttarakhand, Gujarat and Delhi) had more than 50 genomes each, and a total of 1782 genomes were selected from these states. The consortium sequenced sufficiently large samples from each of these states except Delhi and Gujarat. Sequences with more than 10% ambiguous characters were discarded leaving 1696 genomes. CoVa was employed to perform variant analysis on this set of genomes. After generating a Multiple Sequence Alignment (MSA) limited to the sites in the reference genome (NC_045512.2), duplicate genomes were excluded, and further analysis was performed on a final set of 1630 genomes. In this final set, 698 genomes were from the DBT consortium, and the rest were from the other institutes in India.

#### 2. Types distribution

The SARS-CoV-2 genomes selected above were classified into distinct sequence types using CoVa. This sequence typing was based on the nucleotide sequence extracted from 10 positions, proposed to be used for barcoding in Guan *et al.* (16). Sequence types were labeled in the order of their first appearance based on a global dataset of ~17000 sequences collected from GISAID, as of May 21, 2020. Based on this dataset, 17 sequence types were identified and incorporated in the sequence typing program of CoVa. The reference genome belonged to type ST1.

#### 3. Allele frequency dynamics

Allele (mutation) frequency time-series were built for 8 states using collection dates metadata for the selected genomes. The fraction of genomes with a certain mutation of all the genomes that were collected till a certain day were plotted against the number of days passed from the first collection. Only the mutations which were present in at least 1-3rd of the genomes at least once in any state were included in the plot. All the data points from across states were plotted on the same temporal scale *i.e.* the day 0 marked March 01, 2020, the first day of collection in the entire dataset. Further, the earlier data points with less than 20 genomes were excluded from the plot as their frequency estimates would be highly unreliable.

The error in the raw frequency resulting from a small sample size was estimated under the assumption that the raw counts were binomially distributed. For a binomial distribution, the maximum likelihood estimate (MLE) is the same as the raw frequency. Therefore, considering raw frequencies as MLE, the likelihood ratio test can be employed to generate confidence intervals. Briefly, For a significance threshold of **α** = 0.05, a 100(1− **α)** confidence interval can be generated by moving the parameter, binomial probability *p* in this case, away from the MLE to the extent that the null hypothesis of MLE being the true *p* still holds.

#### 4. Binary Clustering

The mutations selected above were clustered based on their presence/absence across 1630 genomes. The binary distance was used to measure the extent of dissimilarity in the occurrence patterns of two mutations. This is the proportion of genomes in which the only one was present out of all the genomes in which either or both of the mutations were present. The mutations were clustered using these pairwise distances in a hierarchical clustering algorithm of complete linkage.

#### 5. Time-series correlation

Mutations were also clustered based on their frequency dynamics. The correlation between any two time-series was estimated using the Spearman’s method, which doesn’t require the relationship between two variables to be linear but only monotonic. The correlation estimates were averaged over the 8 states, and the pairwise correlation matrix was clustered using the same approach as above.

### Haplotype network analysis

For haplotype network, 815 sequences were selected out of the 1034 genome sequences that were generated ad a part of the DBT’s PAN-India 1000 SARS-CoV2 RNA genome sequencing consortium. The genome sequences were aligned to WH01 reference genome using MAFFT (20) with Nextstrain (19) Augur wrapper script. Then the aligned fasta file was converted to Phylip format using a custom Biopython (22) script for haplotype network reconstruction. Using POPART (v 1.7) (23) 717 SARS-CoV-2 genomes were selected with less than 5% undefined states and a TCS Network was created. The network was coloured using the respective residence place (State) of the patient.

## Acknowledgments

Department of Biotechnology, Ministry of Science and Technology, Govt. of India, DBT Wellcome Trust India Alliance Intermediate Fellowship IA/I/16/2/502711 awarded to A.S.N.S., J.C. Fellowship of Department of Science and Technology to SD, Department of Atomic Energy, Government of India, under project no. 12-R&D-TFR-5.04-0800, National Genomics Core at NIBMG and CDFD. NCCS thanks Dr. T. P. Lahane, Director, Department of Medical Education and Research for the permission to use the Maharashtra clinical specimens. The Consortium team would like to thank Dr. Renu Swarup, the Secretary, Department of Biotechnology, Govt. of India, for her enthusiastic support and critical suggestions for this initiative of DBT-AIs.

